# Optimized real time QPCR assays for detection and quantification of *Fusarium* and *Microdochium* species involved in wheat head blight as defined by MIQE guidelines

**DOI:** 10.1101/272534

**Authors:** Elbelt Sonia, Siou Dorothée, Gelisse Sandrine, Cruaud Corinne, Lannou Christian, Lebrun Marc Henri, Laval Valérie

## Abstract

In order to better understand the Fusarium head blight disease, reliable real-time PCR assays for detection and quantification of fungal species belonging to the *Fusarium* and *Microdochium* genus are needed. Specific qPCR assays were developed for nine of those species. All criteria required for reproducing the assays are presented. The assays were species specific and allow quantification of at least 5 pg of fungal DNA and detection of 0.5 pg of fungal DNA per PCR reaction. Moreover we showed that the quantification performances of the tests were not altered in the presence of DNA of closely related species in the sample. The assays were tested on field samples and have been already used in greenhouse experiments.

## 1. Introduction

*Fusarium* head blight (FHB) is a major cereal crop disease causing severe damages worldwide. In addition to reducing yield, FHB leads to the contamination of grains by mycotoxins that are dangerous for human and animal health. FHB involves up to 17 fungal species (Parry et al., 1995) belonging either to the toxinogenic genus *Fusarium* or the non toxinogenic genus *Microdochium*. *F. graminearum F. poae, F. tricinctum, F. avenaceum, F. culmorum M. majus* and *M. nivale* were frequently found in grains showing FHB symptoms. But *F. equiseti, F. acuminatum, F. sambucinum, F. sporotrichioides, F. moniliforme, F. heterosporum, F. subglutinans* and *F. oxysporum* could also be isolated at lower frequency (Ioos et al., 2004). FHB is thus a complex disease and despite intensive investigation, FHB epidemiology as well as the structure and evolution of the species involved in this disease remains poorly understood. A better understanding of the FHB species complex requires accurate, sensitive and specific molecular diagnostic tools, such as real-time PCR (qPCR).

Several parameters influence the reliability of quantification of fungal DNA by qPCR assay (Bustin et al., 2009). First, the DNA extraction protocols from complex samples may vary in their efficiency in removing PCR inhibitors (Cankar et al., 2006). Fredlund et al. (2008) showed that different extraction methods applied to FHB infected wheat resulted in significant differences in the amount of DNA detected. Another study from Brandfass & Karlovsky (2008) showed that increasing the amount of sample material improved the reproducibility of DNA extraction. Finally, Brunner et al. (2009) suggested using a reference gene to normalize the fungal DNA concentration in order to reduce the variation due to DNA extraction. The second main factor that can influence the accuracy of qPCR assay is the reproducibility of the standard calibration curve, which is essential for an accurate DNA quantification. Parameters like slope, y-intercept, and linearity of this curve are thus important. Since detection of early infection is important, qPCR assay should be able to measure small the amounts of fungal DNA. Thus a slight change in the standard calibration curve can greatly alter the measure. For these reasons, the performances of a qPCR assay must be rigorously evaluated in order to ensure the reproducibility of the results. Bustin et al. (2009) proposed a list of essential and desirable information needed to normalize qPCR assays, referred to as the MIQE Guidelines: Minimum Information for Publication of Quantitative Real-Time PCR Experiments.

A major concern in the development of qPCR assays for assessing FHB disease is the specificity of the assay as FHB is caused by species from the same genus. Maybe because the study of FHB has been largely focused on the predominant species *F. graminearum*, few published qPCR methods (Nielsen et al., 2011) take into account the fact that several *Fusarium* species often co-exist in the same samples. The presence of DNA of fungal species from the same genus in the same sample can strongly alter the capacity of the assay to quantify the DNA of a specific species. Using qPCR assays to quantify *F. langesethiae* or *F. graminearum*, Nielsen et al., 2011 showed that the presence of closely related species in the same sample may lead to overestimation of the amount the *F. langesethiae* or *F. graminearum*.

Some papers have already been published with qPCR test for Fusarium species detection (Reischer et al., 2004; Waalwijk et al., 2004; Brandfass & Karlovsky, 2006; Nicolaisen et al., 2009; Demeke et al., 2010). Even more Nicolaisen et al. (2009) have developed qPCR assays for detection of eleven Fusarium species using the single copy gene EF1α, and SYBR Green I fluorescent detection. However the description of performance of each individual assay is sparse and although all the assays could detect their corresponding target in field samples, the ability to quantify the target Fusarium specie in a sample was only perform for *F. graminearum*. Nicolaisen et al., 2009 did not show that their assays using SYBR Green I fluorescent detection, detected and quantified their target with a constant efficiency when other species were present. This might be a concern as Brandfass & Karlovsky (2006) found that SYBR Green I molecules bind at different rates to different amplicons, making quantification less reliable in the case of multiplex analysis. Giglio et al. (2003) showed that SYBR Green I binds preferentially to amplicons with higher GC content and larger size. An interesting alternative would be to develop hydrolysis probes that may be more reliable for diagnosing and quantifying specific species in multi-contaminated samples.

Fewer published papers (Nicholson et al., 1996) present specific PCR tools that are adapted to the species of the genus *Microdochium*. *M. nivale* and *M. majus* have been distinguished as two different species (Glynn et al., 2005) several years ago. These species are regularly found together with *Fusarium* species in wheat grain samples (Xu et al., 2008; Audenaert et al., 2009; Nielsen et al., 2011). Nielsen et al., 2013, used two new q PCR assays for *M. majus* and *M. nivale*. Only the specificity of these assays was described but no other performance criteria. Moreover, the specificity shown was not fully satisfactory as the *M. nivale* assay detected one strain of *M. majus* and the *M. majus* assay detected one *M. nivale* strain out of three.

Our objective in this paper is to enlarge and improve the set of available qPCR assays for FHB species. We present the development of nine qPCR assays specifically designed for the most common species of the FHB complex on wheat in Europe: *F. graminearum, F. culmorum, F. poae, F. avenaceum, F. langsethiae, F. tricinctum, F. sporotrichioides* and the two *Microdochium* species: *M. majus* and *M. nivale*. Following the MIQE guideline (Bustin et al., 2009), the assays were designed to be species specific and to quantify each species in a multi-infected wheat grain. The assays were also assessed for their performance in terms of specificity, reproducibility and ability for quantification of small amounts of DNA in infected samples containing several species.

## 2. Material and methods

The real-time PCR assays were developed using fungal strains from *Fusarium* and *Microdochium* species. Once designed, the assays were assessed for specificity, the reproducibility of the standard curve (R^2^, y-intercept and efficiency). Particularly, the reliability of the quantification was assessed in samples containing several species. Finally, the assays were tested on inoculated or non inoculated field samples.

### 2.1 Material

Fungal strains were isolated from field samples collected at different locations in France in 2008 (Bayer CropScience survey network). The species were identified according to morphological criteria by the Centre de Ressources EQUASA (ESMISAB, 29280 Plouzané - France) and specific end point PCR tests. A subset of strains was selected as representative of the different species of interest and stored on PDA plates at 4°C for DNA extraction and the suspected species name of Fusarium strains was confirmed by sequencing EF1alpha gene.

Infected wheat grains from field plots inoculated with different *Fusarium* species were collected in trials conducted in 2009 and 2010. (Siou et al., 2013). The wheat plants were inoculated either with one or two *Fusarium* species by spraying conidiospores (2.10^4^ spores.mL-1 of each specie) directly on the wheat heads at mid-flowering stage. The samples hereafter referred to as FC, FG, FP, (Table 1; Figure 1) were inoculated by *F. culmorum, F. graminearum* and *F. poae*, respectively. The samples referred to as FG*FC, FG*FP and FC*FP were inoculated with a mixture of *F. graminearum* and *F. culmorum*, or *F. graminearum* and *F. poae*, or *F. culmorum* and *F. poae*, respectively. Two non inoculated fields were also sampled to check for natural infection (samples T1 and T2). After harvest, the seeds were grounded and the flour was stored at room temperature.

**Table 1:** Description of field samples inoculated with *Fusarium* species. Fusarium strains were characterised in Siou (2013).

| Sample name | Wheat variety | Species used for inoculation | Strain name |
| --- | --- | --- | --- |
| FG_1 | Charger | *F. graminearum* | Fg 159 |
| FG_2 | Charger | *F. graminearum* | Fg 178 +Fg159 |
| FG_3 | Trémie | *F. graminearum* | Fg 178 +Fg159 |
| FC_1 | Charger | *F. culmorum* | Fc 129 |
| FC_2 | Charger | *F. culmorum* | Fc 124 |
| FC_3 | Trémie | *F. culmorum* | Fc 124 |
| FC_4 | Trémie | *F. culmorum* | Fc 285 |
| FP_1 | Charger | *F. poae* | Fp3 |
| FP_2 | Trémie | *F. poae* | Fp3 |
| FG*FC_1 | Charger | *F. graminearum* * *F. culmorum* | Fg 159 * Fc 129 |
| FG*FC_2 | Charger | *F. graminearum* * *F. culmorum* | (Fg 178 +Fg159)* Fc 129 |
| FG*FC_3 | Charger | *F. graminearum* * *F. culmorum* | Fg 159 * Fc 124 |
| FG*FC_4 | Charger | *F. graminearum* * *F. culmorum* | (Fg 178 +Fg159)* Fc 124 |
| FG*FC_5 | Trémie | *F. graminearum* * *F. culmorum* | (Fg 178 +Fg159)* Fc 129 |
| FG*FC_6 | Trémie | *F. graminearum* * *F. culmorum* | (Fg 178 +Fg159)* Fc 124 |
| FG*FP_1 | Charger | *F. graminearum* * *F. poae* | Fg 159 * F3 |
| FG*FP_2 | Charger | *F. graminearum* * *F. poae* | (Fg 178 +Fg159)* F3 |
| FG*FP_3 | Trémie | *F. graminearum* * *F. poae* | (Fg 178 +Fg159)* Fp3 |
| FC*FP_1 | Charger | *F. graminearum* * *F. poae* | Fc 124 * Fp3 |
| FC*FP_2 | Trémie | *F. graminearum* * *F. poae* | Fc 124 * Fp3 |
| T1 | Charger | No inoculum |  |
| T2 | Trémie | No inoculum |  |

**Figure 1:**
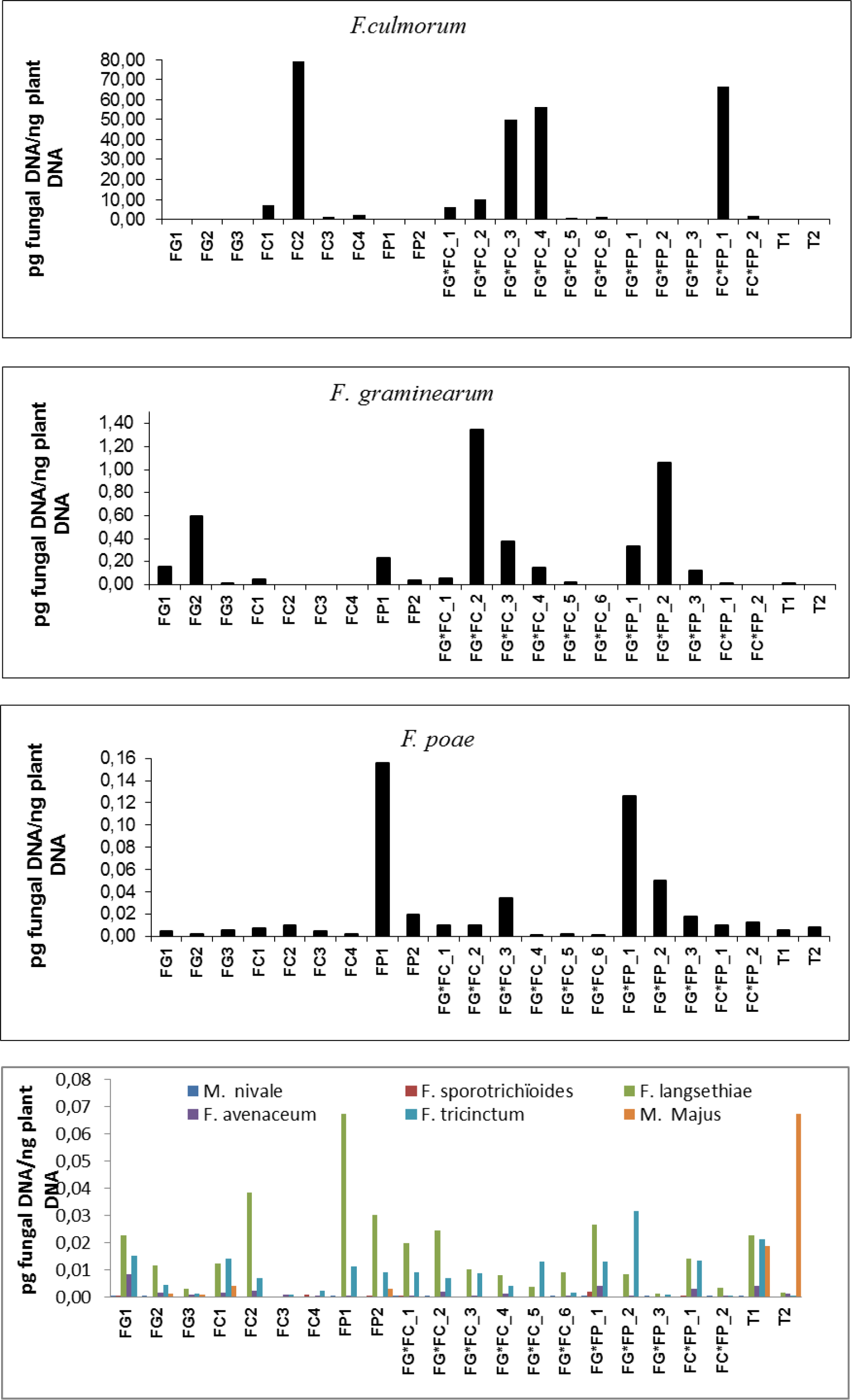
Amounts of fungal DNA measured by qPCR species specific assay in samples from inoculated or non incoculated field plots. **a**: quantification of *F. culmorum*, **b**: quantification of *F. graminearum*, **c**: quantification of *F. Poae*, **d**: quantification *F. langsthetiae*, *F.sporotrichoides, F. tricinctum, F. avenaceum*, *M. nivale, M. majus*. Plots were inoculated with one species *F. culmorum* (FC); *F. graminearum* (FG); *F. poae* (FP) or with two species *F. graminearum* and *F. culmorum* (FG*FC); *F. graminearum* and *F. poae* (FG*FP), *F. culmorum* and *F. poae* (FC*FP). T1 and T2 were sampled from non inoculated plots. The quantities are expressed in pg of fungal DNA per ng of plant DNA in each sample.

### 2.2 DNA extraction

Two methods (A and B) were used for DNA isolation from the strains cultured on artificial medium. For method A, the isolates were grown on PDA agar plate. For method B, the isolates were grown on PDA plates covered with a cellulose film (Octaframe, Copenhagen, Denmark). In both cases, the fungus was grown for 2 weeks at 23°C with a 12- hours light period. For method A, the mycelium was scrapped directly from the PDA plate with a spatula and weighted, then 200 mg of mycelium was used for DNA extraction using DNeasy Plant Mini kit (Qiagen, Courtabeuf). The extraction was performed according to the manufacturer’s instructions with only minor modification: incubation time at 65°C was prolonged for one hour and the volume of AP1 and AP2 buffers was doubled. For method B, the mycelium was scrapped from the cellulose film and lyophilised for 24h in a Edwards Modulyo 4K Lyophilizer (Edwards, United Kingdom). The DNA was extracted as in method A except that, after mixing the AP2 buffer, 3 successive centrifugations were performed on the supernatant until it was completely limped. The DNA quality and quantity were determined using a NanoDrop^®^ ND-1000 spectrophotometer (Labtech, Paris).

A third extraction method was used for infected wheat field samples. The grains collected from the field trial were grounded in a mixer mill MM 400, RetschR156 (Retsch, France) for 1 min 30 sec at 30 Hz. A sample of 100 mg of this powder was used for DNA extraction using the DNeasy Plant Mini kit (Qiagen, Courtabeuf) according to the manufacturer’s instructions. Only minor modification were applied: the samples were sonicated in the AP1 buffer for 2 min at 42Khtz, then Proteinase K (20 mg.ml-1 final concentration) was added to this mix; the samples were incubated for 1 hours at 65°C; the AP1 and AP2 buffers volume were doubled. The DNA quality and quantity were determined using a NanoDrop^®^ ND-1000 spectrophotometer (Labtech, Paris).

### 2.3 DNA sequencing to confirm strain characterization

For further characterization of *Fusarium stains*, the *EF1 α* region was amplified using EF1-Fusa-spp-Bioger-F 5’ TCGTCGTCATCGGCC 3’ and EF1-Fusa-spp-Bioger-R 5’AGTGATCATGTTCTTGATGAAATC 3’. The PCR reaction contained 1 × mastermix for the Taq DNA pol EPRQC200 (MP biomedical, France), 5mM dNTPs, and 0.3 μM of each primer in a total volume of 40 μL. PCR reactions were performed with either 30 ng of genomic DNA. PCR reactions were performed using an Applied Biosystems 9700 thermocycler. The amplification parameters consisted of an initial denaturation step at 95°C for 5 min; then either 40 cycles of denaturation at 95°C for 30 s, annealing at 54°C for 30 s, and elongation at 72°C for 60 s; and a final extension step at 72°C for 10 min. After checking for the expected size of the fragment on an agarose gel, the PCR fragments were sent to sequencing using forward and reverse primers to the Genoscope (Centre National de Séquençage, Evry, France) in the frame of the project@SPEED-ID. After careful chromatogram check, sequences were deposit in R-Syst::fungi database (http://138.102.89.206/new_rsyst_chp/; see supplementary S1 for sequence accession number).

### 2.4 Real-time PCR set up

*EF1α* gene sequences from the 7 targeted *Fusarium* species were aligned using MultAlin [31] (Corpet, 1988) to find species specific primers. Several sets of candidate primers were designed were designed using the Primer ExpressTM software, version 2.0 (Applied Biosystems, Foster City, CA, USA). Their specificity was first assessed *in silico* by BLAST with the NCBI database. They were then synthetized by Eurogentec (Seraing, Belgium).

For *Micordochium majus* and *nivale*, another monocopy gene the β tubulin gene was choosen for the development of the qPCR assay. The same procedure as for Fusarium (PCR, sequencing) was performed with for *M. nivale* and *M. majus* species. The primer used for amplicafication and sequencing of a fragment of β tubulin were the following one’s *BtubfusaterF* 5’ CACGGTCTCGACAGCAATG 3’, BtubfusaTerR 5’ATGGTACCGGGCTCGAGAT 3’. Several sets of candidate primers for the q PCR assays were designed were designed using the Primer ExpressTM software, version 2.0 (Applied Biosystems, Foster City, CA, USA). Their specificity was assessed using several strains of *.majus* and *M nivale* and in silico on other species.

The primers and probes selected and assessed in this study are listed in Table 2. All the probes used were Taqman^®^ labelled with the FAM/TAMRA reporter/quencher dye.

**Table 2:**
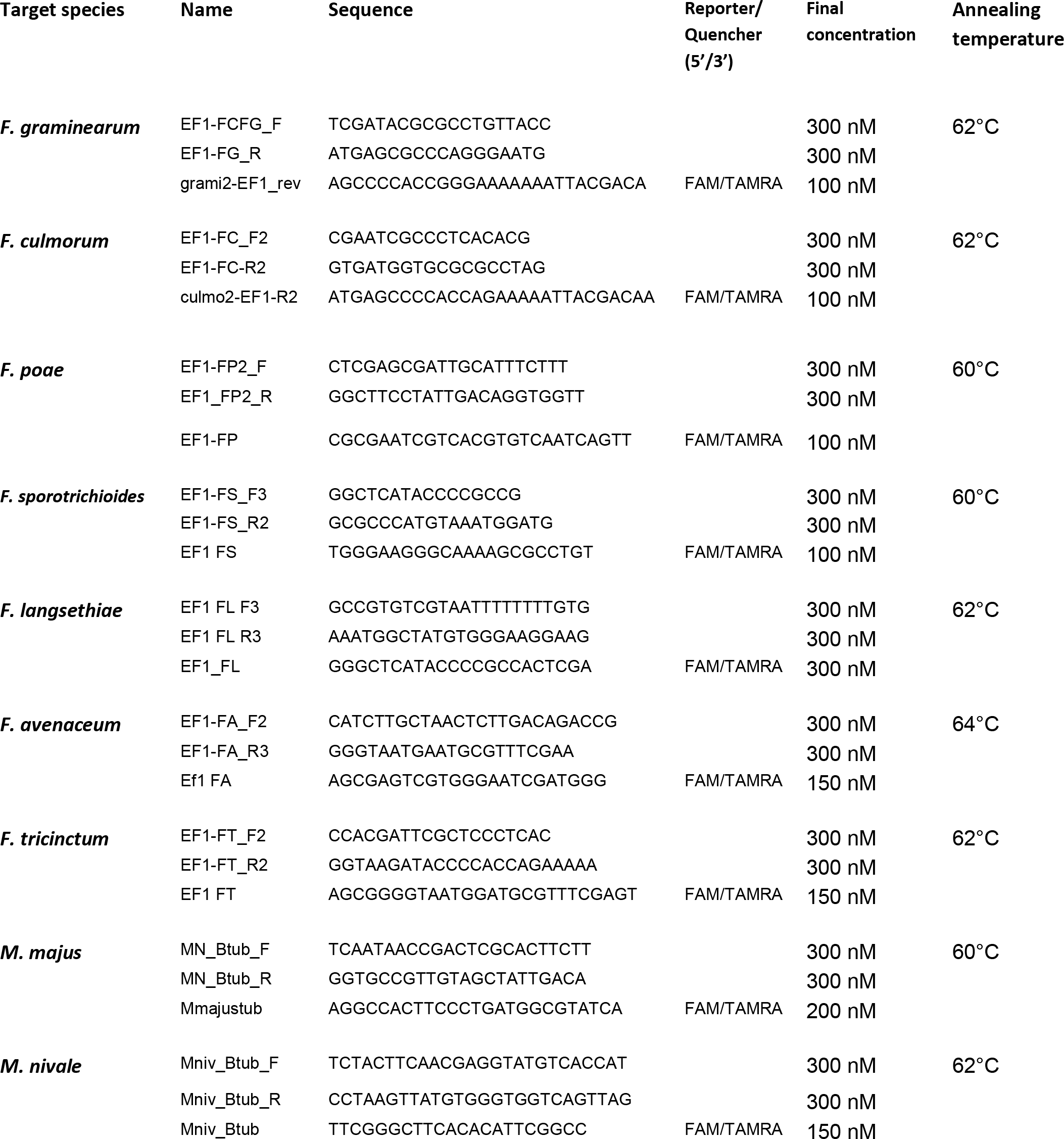
Primer and probe sequences and qPCR amplification conditions for each *Fusarium* and *Microdochium* species.

All the qPCR assays developed in this study were performed using the 2X qPCR MasterMix with ROX and Uracil N Glycosylase (UNG) from Eurogentec (Angers, France). The PCR reactions were performed in a total of 25μl consisting of 12.5 μL of mastermix, with primers and probes at the concentrations described in Table 2. For standard calibration curve, reference DNA concentrations ranged from 5 ng to 0.005 ng. In each experiment, DNA were analysed in triplicates. A standard curve was performed on each PCR plate. The real-time PCR reactions were performed on an ABI PRISM 7900 Sequence Detection System (Applied Biosystems, Foster City, CA, USA) in Applied Biosystem 96 well plates. Cycling conditions were an initial denaturation step of 95°C for 10 min followed by 40 cycles each consisting of 95 °C for 15s then the temperatures specified in Table 2 for one min.

For wheat sample quantification, a plant reporter gene was used to normalize the fungal amount per sample. The plant DNA was quantified in the samples from the field trials using plant EF1 qPCR primers, as described in Nicolaisen et al. (2009).

### 2.5 Performance analyses of the qPCR assay

The species specificity of each primer/probe set was tested by quantifying 10 ng of DNA of at least three strains of the following *Fusarium* species: *F. sporotrichioides, F. tricinctum, F. langsethiae, F. graminearum, F. avenaceum, F. poae, F. culmorum, M. nivale, M. majus* and two strains of *F. equiseti, F. moniliforme, and F. subglutinans*.

Linearity, efficiency and y-intercept, were estimated from standard curves performed with 10- time serial dilutions of fungal genomic DNA (from 5 ng to 0.005 ng). The slope of the standard curve was used to calculate the qPCR efficiency with the following formula: PCR efficiency=10-1/slope-1 (Bustin et al., 2009). The y-intercept was calculated from the standard curve formula: log(quantity)= slope *(Cq value) + y-intercept. The linearity was estimated by the R^2^ of the standard curve. A minimum of three standard curves established with different strains of each species was used to estimat the efficiency, linearity and y-intercept values for each assay (table S1). The influence of the DNA extraction method (A or B) on the standard calibration curve and the performance criteria was tested for *F. graminearum* only.

The influence of the presence of a closely related species in the same DNA sample was assessed using a ten time serial dilution of the targeted DNA from 5ng to 0.005ng per PCR reaction. Mixed samples corresponding to pooled DNAs from different *Fusarium* species at 1 ng/μl (Table 6) were added to these serial dilutions before performing real-time PCR experiments.

The final criteria to be tested were the limit of detection (LOD) and the limit of quantification (LOQ). To calculate for the LOD and LOQ, ten times serial dilutions were performed ranging from 5ng to 0.0005ng and two intermediates point of respectively 0.001ng and 0.0001ng are added. For the LOD, the last dilutions 0.0001ng and 0.0005ng were estimated using 20 replicates from the same experiment. By convention, the LOD was reached when 5% of the replicates were negative (one out of 20). For the LOQ, we used 10 replicates of the tested dilution to estimate the mean Cq value. The LOQ Cq value was reached when the standard deviation of the mean Cq value of 10 replicates was lower than 0.5 Cq values.

The influence of intraspecific variation among strains from the same species on DNA quantification was tested for each assay. To this end, genomic DNA was extracted from a minimum of three from the same species and 1 et 10 ng of each DNA was requantified by the corresponding qPCR assay (table S2).

### 2.6 Validation of the qPCR assays on field samples

As described by Nicolaisen et al. (2009), the amount of fungal DNA was calculated from Cq values using the standard curves, and these values were normalized with the estimated amount of plant DNA of the same sample, quantified by wheat-specific qPCR assay. The quantification of the plant DNA using plant EF1 qPCR primers and SYBR Green I were performed as described in Nicolaisen et al. (2009). For the plant EF1-based assay, qPCR was carried out in a total volume of 25μl, consisting of 12.5μl of MESA GREEN qPCR MasterMix Plus for SYBR^®^ Assay (Eurogentec Angers, France), 300nM of each primers and 5μl of DNA at 30ng/μL. The cycling conditions were an initial denaturation step at 95°C for 10 min followed by 40 cycles, each consisting of 95 °C for 15s and 60°C for 1 min followed by dissociation curve analysis at 60 to 95°C. The results were analysed with the AB SDS2.2.2 software (Applied Biosystem). In each experiment, DNA was analysed in triplicates and a standard curve was performed on each PCR plate.

## 3. Results

### 3.1 Assay design

In the course of the development of our assays Nicolaisen et al. (2009) published 11 qPCR assays for FHB detection. We decided to use their assays for *F. sporotrichioides* and *F. langsethiae* detection since our own assays for those species were not ready. However we encountered transferability problems may be due to a difference in equipment or products between the two laboratories (data not show) and we then decided to complete the development of our assays using hydrolysis probes with the objective to improve their specificity. We thus developed new assays for *F. sporotrichioides, F. tricinctum, F. langsethiae, F. graminearum, F. avenaceum, F. poae, F. culmorum* and for *M. nivale*, and *M. majus*. Inter-and intra-specific sequence variability was analysed in silico for the EF1α gene for *Fusarium* and for the β tubulin gene for *Microdochium*.

### 3.2 Performance analyses

Following Bustin et al (2009), we evaluated the specificity, the efficiency, the y-intercept, the linearity, the limit of detection and the limit of quantification of each assay.

For the specificity assessment, we tested specificity of our qPCR assays against different Fusarium species and the 2 Microdochium ones. The specificity of Microdochium test was also assessed on Fusarium species. The Fusarium and Microdochim strains used for specificity analysis were first characterized by morphological description when possible. Their taxonomic name was then confirmed by sequencing EF1 alpha gene. The mean Cq values obtained when quantifying DNA from at least 3 strains of each species are presented in Table 3. In all the tests, we only detected the targeted species, except for the *F. culmorum* and *F. cerealis* which remained undistinguishable as reported by Nicolaisen et al. (2009). The *F. tricintum* assay also slightly detected *F. avenaceum* when the annealing/elongation temperature was lower than 62°C (Cq : 39). However this phenomenon did not occur when increasing the annealing/elongation temperature to 62°C. For each species, the Cq values obtained for DNA quantification from three different strains were similar as the standard deviation of the mean calculated from these Cq values was lower than 0.5Cq. As the assays were specific for Fusaium species we did not check other wheat fungal pathogens which are less closely related.The performance of these assays was estimated using values computed from standard curves such as efficiency, linearity and y-intercept (Table 4 and detailed in S1). Each assay showed efficiency above the 90% preconised by GMO guidelines and was reproducible, with a standard deviation lower than 3.5%. The linearity was respected over at least 4 range of magnitude and was not affected by two additional dilutions (data not shown). The y-intercepts were reproducible with a standard deviation lower than 0.5.

**Table 3:**
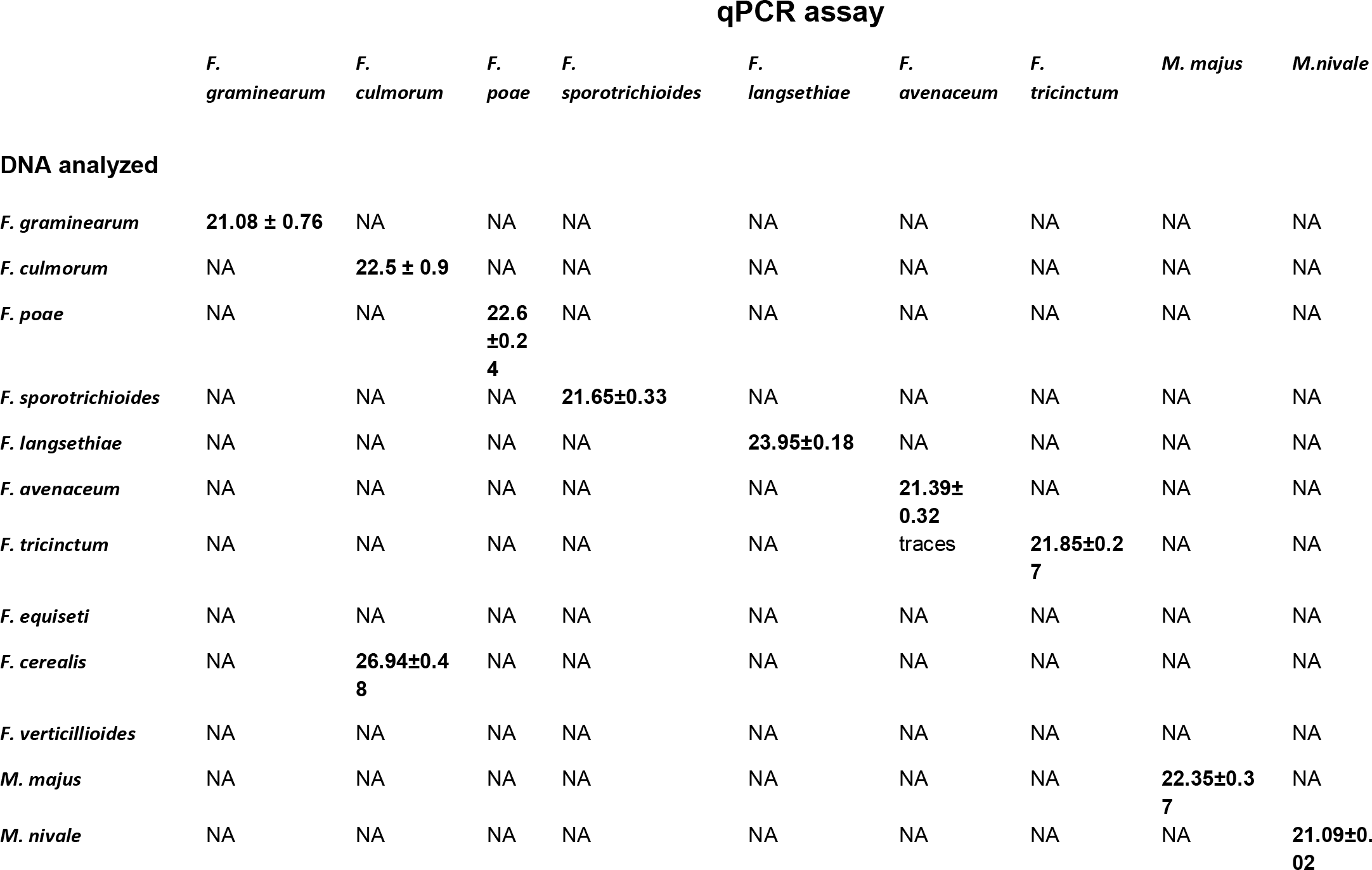
Analysis of the specificity of *Fusarium* and *Microdochium* qPCR assays. Cq value and standard deviation obtained when 10 ng of DNA from 3 strains per species was quantified independently by qPCR. NA: no amplification

**Table 4:** Performance of *Fusarium* species-specific qPCR assays. For each assay, efficiency, *y*-intercept and linearity were calculated from standard curves established with DNA extracted from at least 3 strains

| Target species | Efficiency<sup>a</sup> | y- intercept<sup>b</sup> | Linearity (r<sup>2</sup>) |
| --- | --- | --- | --- |
| *F. graminearum* | 94.29 ± 2.00 | 24.92 ± 0.35 | 0.998 ± 0.001 |
| *F. culmorum* | 98.06 ± 2.22 | 25.33 ± 0.14 | 0.996 ± 0.002 |
| *F. poae* | 94.8 ± 0.94 | 25.01 ± 0.1 | 0.997 ± 0.0006 |
| *F. sporotrichioides* | 96.87 ± 3 | 24.6 ± 0.27 | 0.998 ± -0.01 |
| *F. langsethiae* | 95.43 ± 2.5 | 26.25 ± 0.16 | 0.997 ± 0.002 |
| *F. avenaceum* | 96.68 ± 3.38 | 25.43 ± 0.48 | 0.996 ± 0.002 |
| *F. tricinctum* | 98.66 ± 0.23 | 25.27 ± 0.3 | 0.997 ± 0.0006 |
| *M. majus* | 96.07 ± 3.2 | 25.72 ± 0.38 | 0.995 ± 0.001 |
| *M. nivale* | 97.1 ± 1.2 | 24.02 ± 0.67 | 0.998 ± 0.002 |

a: efficiency = $10^{-1/\text{slope}} - 1$
b: y-intercept is calculated from the standard curve as $\log(\text{quantity}) = \text{slope} (\text{Cq value}) + \text{y-intercept}$

All the assays reached a limit of detection (LOD) of 0.5 pg of fungal DNA in the PCR reaction. With an additional dilution (0.1 pg), amplification was however observed in some replicates. All the assays had a limit of quantification (LOQ) of 5 pg of fungal DNA per PCR reaction.

We then check the quantification accuracy of the assays on genomic DNA extracted from three strains of the same species. The DNA was quantified by NanoDrop spectrophotometer and by qPCR (Table 5 and detailed in S2). Similar results were obtained with both methods, showing that our qPCR assays accurately quantified fungal DNA and that strain had no influence on quantification.

**Table 5:**
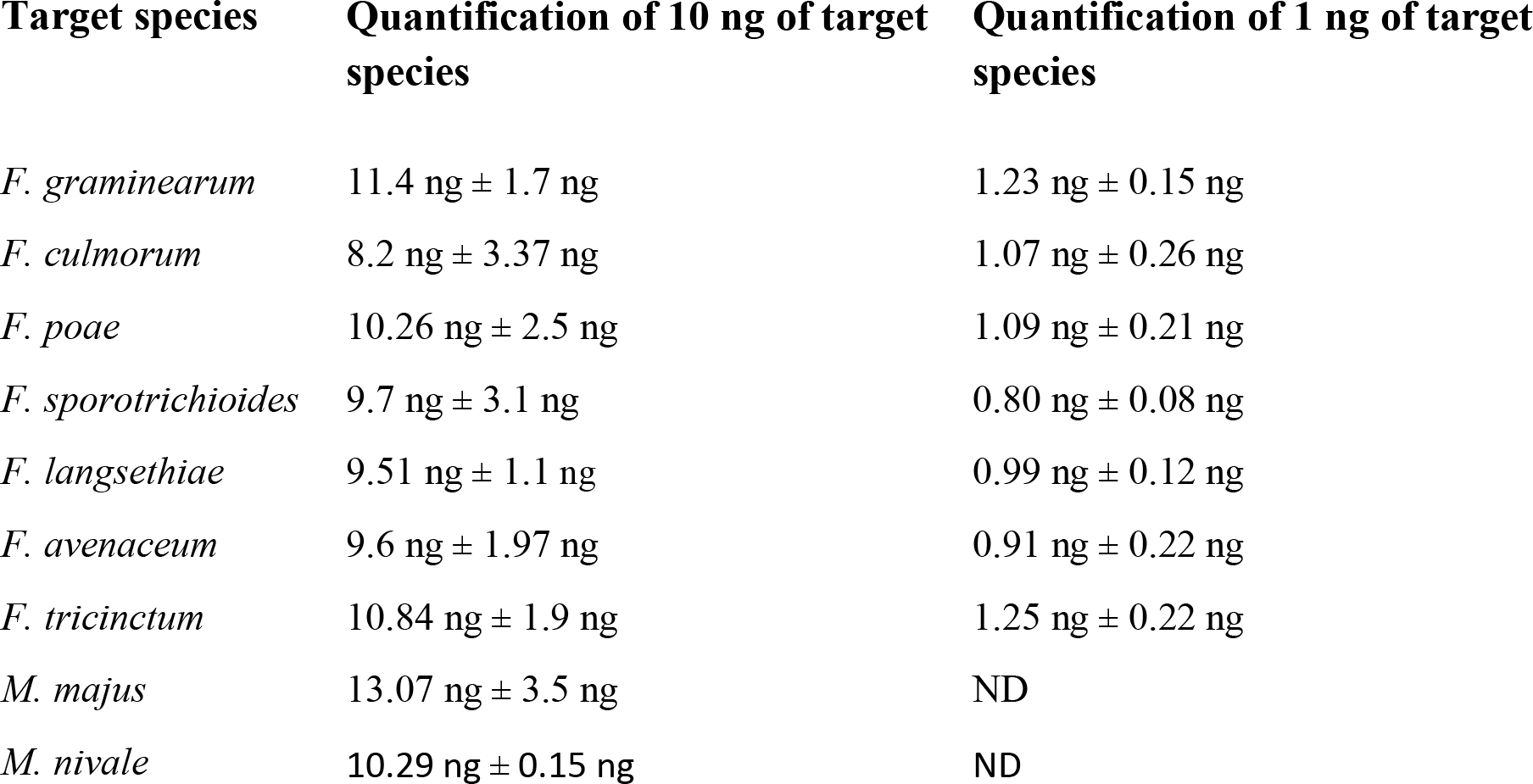
Mean and standard deviation of qPCR quantification of 10 ng and 1 ng of fungal DNA from 3 strains for each species.

In order to check for the reproducibility of the assays in complex samples, the performance (the efficiency, the y-intercept, the linearity) of our qPCR assays were checked again in the presence of DNA from closely related species. No significant influence of was noticed on performance criteria of the assays (data not shown). The accuracy of the quantification was also evaluated in the presence of DNA mixture. Similar values to the one obtained with target DNA alone were obtained in this experiment (Table 6). There is no interference in the quantification of the target species by other DNA from related species (Table 6). For *F. culmorum* and *F. poae*, an inhibitory effect was observed at the highest concentration of the target DNA (5 ng), but this effect was not observed at lower DNA concentrations. For all assays, the quantified values were close to the expected values even for low amounts of the target DNA (0.005 ng).

**Table 6:** Quantification and specificity of *Fusarium* species-specific qPCR following spiking with a mixture of DNA preparations from different species.

| Target species | Other species present in the mixture | Quantification of 5 ng of target species | Quantification of 0.5 ng of target species | Quantification of 0.05 ng of target species | Quantification of 0.005 ng of target species |
| --- | --- | --- | --- | --- | --- |
| ***F. graminearum*** | *F. culmorum*; *F. langsethia*; *F. tricinctum*; *F. poae*; *F. avenaceum*; *F. sporotrichoides*; *M. nivale* ; *M. majus* ; *F. equiseti*; *F. verticillioides* | 4.88 ± 0.50 | 0.55 ± 0.05 | 0.055 ± 0.004 | 0.0055 ± 0.0006 |
| ***F. culmorum*** | *F. graminearum*; *F. langsethia*; *F. tricinctum*; *F. poae*; *F. avenaceum*; *F. sporotrichoides*; *M. nivale* ; *M. majus* ; *F. equiseti*; *F. verticillioides*; *F. subglutinans* | 3.67 ± 0.76 | 0.46 ± 0.04 | 0.046 ± 0.004 | 0.0045 ± 0.0018 |
| ***F. poae*** | *F. graminearum* *F. culmorum*; *F. langsethia*; *F. tricinctum*; *F. avenaceum*; *F. sporotrichoides*; *M. nivale* ; *M. majus* ; *F. equiseti*; *F. verticillioides*; *F. subglutinans* | 3.46 ± 0.03 | 0.38 ± 0.01 | 0.042 ± 0.008 | 0.0052 ± 0.0009 |
| ***F. sporotrichoides*** | *F. graminearum* *F. culmorum*; *F. langsethia*; *F. tricinctum*; *F. avenaceum*; *F. poae*; *M. nivale* ; *M. majus* ; *F. equiseti*; *F. verticillioides*; *F. subglutinans* | 3.9 ± 0.55 | 0.44 ± 0.025 | 0.046 ± 0.004 | 0.004 ± 0.0006 |
| ***F. langsethia*** | *F. graminearum* *F. culmorum*; *F. tricinctum*; *F. avenaceum* *F. sporotrichoides*; *F. poae*; *M. nivale* ; *M. majus* ; *F. equiseti*; *F. verticillioides*; *F. subglutinans*, *F. cerealis* | 4.85 ± 0.99 | 0.50 ± 0.12 | 0.054 ± 0.001 | 0.0047 ± 0.0002 |
| ***F. aveaceum*** | *F. graminearum* *F. culmorum*; *F. langsethia*; *F. tricinctum*; *F. sporotrichoides*; *F. poae*; *M. nivale* ; *M. majus* ; *F. equiseti*; *F. verticillioides*; *F. subglutinans* | 4.75 ± 0.07 | 0.51 ± 0.03 | 0.052 ± 0.004 | 0.005 ± 0.0005 |
| ***F. tricinctum*** | *F. graminearum* *F. culmorum*; *F. langsethia*; *F. avenaceum*; *F. sporotrichoides*; *F. poae*; *M. nivale* ; *M. majus* ; *F. equiseti*; *F. verticillioides*; *F. subglutinans* | 4.46 ± 0.39 | 0.52 ± 0.14 | 0.042 ± 0.002 | 0.0045 ± 0.0002 |
| ***M. majus*** | *F. graminearum* *F. culmorum*; *F. langsethia*; *F. tricinctum*; *F. avenaceum*; *F. poae*; *M. nivale* ; *F. equiseti*; *F. verticillioides*; *F. subglutinans*; *F. cerealis* | 3.14 ± 0.65 | 0.4 ± 0.05 | 0.0415 ± 0.09 | 0.0037 ± 0.0009 |
| ***M. nivale*<sup>a</sup>**<br>**(10 ng and dilutions)** | *F. graminearum* *F. culmorum*; *F. langsethia*; *F. tricinctum*; *F. avenaceum*; *F. poae*; *M. majus* ; *F. equiseti*; *F. verticillioides*; *F. subglutinans*; *F. cerealis* | 9.6 ± 0.70 | 0.97 ± 0.11 | 0.082 ± 0.001 | 0.0077 ± 0.0010 |

<sup>a</sup> for *M. nivale*, dilutions to between 1 and 10 ng DNA per PCR were analyzed

As far as we are aware this is the first report in which DNA quantification is done with mixture of DNA from related fungal species.

### 3.3 Detection of FHB species in infected wheat grains

Table 1 describes the sample used for this analysis. DNA was extracted in a reproducible way for all samples from 100mg of grounded flower. The mean concentration value obtained for all sample was 139 +/-27 ng/μl in a total volume of 100μl. Samples from inoculated and non inoculated field plots were tested with each of the assay presented in this paper. The results are presented in Figure 1. High amount of *F. culmorum* were detected in inoculated plots (up to 80 pg fungal DNA / ng plant DNA in sample FC2) but not in non-inoculated plots T1 and T2, Less than 0.2 pg fungal DNA /ng plant DNA were quantified in T1 or T2 likely resulting from natural contamination (Figure 1A). The levels of contaminations were lower for *F. graminearum* (Figure 1B). Even in inoculated plots, the highest amount of *F. graminearum* found was only 1.5 pg/ng in FG*FC2 sample. Inoculation of *F. poae* did not work well since the level of contamination was found at a maximum of 0.15 pg fungal DNA / ng plant DNA in the FP2 sample (Figure 1C). This is consistent with the disease severity values based on visual observations found by Siou, (2013). In the plots inoculated with two FHB species, the two species were quantified. For example, in sample FG*FP1, the amount of *F. graminearum* and *F. poae* DNAs were respectively 0.33 and 0.12 pg fungal DNA/ng of plant DNA (Figure 1). Samples T1 and T2 (non inoculated field plots) were used to assess natural contaminations. All the species except *F. sporotrichoides* and *M. nivale* were quantified in at least one of these two plots (Figure 1D). In fact, we were able to quantify the natural contamination in inoculated plots as well. *F. langsethiae* and *F. tricinctum*, were the most frequent contaminant species. *F. sporotrichioides* was detected in only 5 samples out of 24 but at very small amounts (<0.0018 pg/ng of plant DNA). *M. nivale* was present in 8 samples out of 24 at a very low level (<0.0003pg/ng).

## 4. Discussion

The relative frequency of the FHB species is often measured in epidemiological and population studies (e.g.; Parry et al., 1995; Xu et al., 2005; Chandelier et al. 2011; Siou et al., 2013). Classical methods rely on microbiological isolation and morphological criteria that are not always sufficient to identify these species accurately. Real-time quantitative PCR (qPCR) is the most accurate and efficient way to detect species in environmental samples. We have developed nine qPCR assays for detection and quantification of *Fusarium* and *Microdochium* species responsible for FHB in France and in Europe: *F. graminearum, F. poae, F. tricinctum, F. avenaceum*, as well as some less frequent species, *F. culmorum, F. sporotrichioides, F. langsethiae*, (Ioos et al., 2004; Xu et Nicholson, 2009). We have chosen the monocopy gene EF1 α to develop the assays for *Fusarium* species. This gene is efficient to distinguish related *Fusarium* species (O’ Donnell et al., 2008; Nitschke et al., 2009) and used for the FUSARIUM-ID database (Geiser et al., 2004). Sequences of FHB *Fusarium* species were downloaded from NCBI and the FUSARIUM-ID V. 1.0.I databases and aligned, confirming that this gene provides sufficient inter-species variability to distinguish related species while intra-species variability is minimal. The species *M. majus* and *M. nivale* are also involved in FHB and sometimes found as a dominant species (Nielsen et al., 2011). As far as we know, only one publication (Nielsen et al., 2013) proposes qPCR assays that differentiate the two species. But at that time we already had analyzed the potential of our two new qPCR assays using the β tubulin gene for specifically detecting *M. majus* and *M. nivale* in samples contaminated by *Fusarium* species.

Problems of transferability between laboratories are important issues regarding qPCR assays. A work proposing similar assays was published during the course of our work (Nicolaisen et al., 2009). As our own assays for *F. sporotrichioides* and *F. langsethiae* were not fully developed. We attempted to transfer their assays in our laboratory) Although we used the same primer and probe concentrations, the same PCR conditions and the same qPCR apparatus, the qPCR reactions were realized with a qPCR mix obtained from a different company. In these conditions, their test did not show a good performance (lack of specificity, efficiencies problems). We decided to continue our assay to get a set of qPCR test developed in the same wayfor FHB epidemic studies.

For qPCR assays, another important point is the specificity. This is particularly important for FHB species since these fungi are closely related and frequently present in the same field or grain sample (Audenaert et al., 2009). Two technologies are currently used for qPCR: SYBR Green I and hydrolysis probes. To improve the specificity of the assays proposed, we used hydrolysis probes, which allows differentiating sequences with only two base differences (Livak., 1999). With this technique, eight assays out of nine proved species specific. The *F. culmorum* assay was able to detect and quantify at the same level *F. cerealis* DNA. This is actually not surprising since only one base was found different between the sequences of *F. culmorum* and *F. cerealis*. So when using our qPCR assay to detect *F. culmorum*, we advise to use another assay specifically designed to detect *F. cerealis* in order to make sure which species is present. A PCR specific assay for *F. cerealis* has been published by Fernández-Ortuño et al. (2010) and it could be used in complement to our assays.

In order to produce reproducible and transferable protocols, the assays developed in this study followed the rules proposed by Bustin et al. (2009). The document for GMO testing in food and feed is also a reference for developing precise and reliable assay for DNA quantification by qPCR (Definition of Minimum Performance Requirements for analytical methods of GMO testing, October 2008). This document recommends that the standard curve established for each assay should be linear over at least four (and up to five) ranges of dilution, with an efficiency above 90% and coefficient of correlation above 0.99. These criteria were successfully met for each assay on experiments repeated independently at least three times. The stability of the y-intercept of the standard curves was also evaluated as it has a major impact on DNA quantification. In fact, having similar efficiency values but different y-intercept changes the quantities of DNA obtained for a sample. This criterion is rarely evaluated in publications presenting assays for *Fusarium* detection and quantification. It was not described for *Fusarium* assays in Nicolaisen et al. (2009) nor for *Microdochium* assays proposed by Nielsen et al., 2013. The assays developed in this study showed reproducible y-intercept values which should insure reproducible quantification.

The quantification performances of our assays were evaluated for the targeted species, either alone or in the presence of other species. The presence of DNA from other species in the same sample could impair the quantification capacity of the assay. Quantification of a 10- fold dilution, from 5 to 0.005ng, was not altered by the presence of the DNA from closely related species. As far as we are aware, such quantification in the presence of closely related DNA has rarely been performed for FHB species. Nielsen et al. (2011), also evaluated the quantification capacity of their qPCR assays in the presence of another species, but they only checked the performance of the assay in a background of the closest related species i.e.: *F. culmorum* DNA for the *F. graminearum* assay. They noticed an increase in the efficiency of the qPCR assays which could induce overestimation of the amount of the target DNA. In our approach, we mixed several species including the most closely related ones, and this did not alter the performances of the different assays. Even though adding DNA of several strains in the mix artificially accumulates PCR inhibitors, this did not impair the assay performances.

The performance of qPCR amplification is influenced by different factors like the presence of inhibitors co-purified with DNA during extraction, primer and probe concentrations and the mix used for the PCR reaction. The quality of the DNA used for the standard curve is crucial for quantification by qPCR (Fredlund et al., 2008). A poor DNA quality may interfere greatly with NanoDrop spectrophotometer quantification and PCR amplification. Such effects were observed in our study. Even if there was little differences between the extraction methods A and B, the reproducibility of the standard curve was noticeably improved when fungal DNA was extracted with method B and this was true for each assay. Potential inhibitors had an impact on the reproducibility of two parameters that are essential for DNA quantification: the y-intercept and the PCR efficiency (data not shown).

Our assays were used to analyse samples from a field trial inoculated with known FHB species. As proposed by Brunner et al. (2009), we choose to express the quantity of fungal DNA relative to a reference plant gene. This avoids a bias due to differences in the efficiency of the DNA extraction. The inoculated species were detected even when the targeted species was co-inoculated with other species. Quantification revealed differences in infection levels showing that some inoculations did not work properly. This was consistent with the visual observation of the symptoms (Siou, 2013). Natural contaminations that normally occur in the fields were also detected.

Our assay *F. graminearum*, *F. culmorum* and *F. poae* assays were also used by Siou et al., (2013) to correlate fungal DNA with visual symptoms and toxin content on wheat head after inoculation of either single species or species mixtures. In all these experiments, the toxin-DNA relation was linear and highly significant. The same tests were used in another study to detect fungal DNA from FHB grains inoculated either alone or in mixtures on individual wheat spikes. These experiments showed clear exclusion of the FHB species at the spike level and the data obtained confirms both the specificity and the precision of our assays (Siou, 2013).

## 5. Conclusion

We propose nine qPCR assays specifically designed for the most common species of the FHB complex on wheat in Europe: *F. graminearum, F. culmorum, F. poae, F. avenaceum, F. langsethiae, F. tricinctum, F. sporotrichioides* and the two *Microdochium* species: *M. majus* and *M. nivale*. The assays were carefully assessed according to MIQE guideline and described in details so that they could be used by over laboratories for detection of *Fusarium* and *Microdochium* species in wheat FHB. The assays presented here are adapted to studies FHB field samples, as the presence of related species does not modify the detection and quantification performances of the tests. It is to be noticed that a poster presenting the transferability of our assays was presented at the 12th European Fusarium Seminar in 2013 in Bordeaux. The qPCR tests proved to be transferable as the only significant effect on result variation was DNA extraction.

## Acknowledgements

The authors are grateful to Bayer CropScience and Arvalis Institut du Végétal for their financial support. This work was financially supported by the French ANR Project DON&Co 10-CESA-01.

This work was part of the project@SPEED-ID “ Accurate SPEciES Delimitation and IDentification of eukaryotic biodiversity using DNA markers” proposed by F-Bol, the French Barcode of life initiative.

